# Using peptide-exchange systems to interrogate peptide-specific KIR binding to HLA Class I

**DOI:** 10.64898/2026.03.03.708729

**Authors:** Tanusya M. Murali, Beining Li, Emery Hoos, Lucy Collinson, Eric O. Long, Omer Dushek, Tim Elliott, Malcolm J. W. Sim

## Abstract

The killer-cell immunoglobulin-like receptors (KIR) are a family of activating and inhibitory HLA class I (HLA-I) binding receptors expressed on natural killer (NK) cells and subsets of T cells. The KIR detect HLA-I molecules in a peptide-dependent manner, with some KIR displaying exquisite peptide-specificity. Studying peptide recognition by KIR often uses TAP-deficient cell lines expressing single HLA-I alleles, which are heterogenous and time consuming to generate. Here, we established an alternative approach using peptide-exchange technologies hitherto developed for studying T cell recognition of HLA-I. We tested two methods; dipeptide-mediated peptide exchange and ‘open-HLA-I’, HLA-I molecules consisting of heavy chain-β_2_m disulphide bonded dimers. We combined peptide-exchange technologies with SpyTag-SpyCatcher chemistry to allow rapid detection of KIR binding via HLA-I displayed on plates or cells. We demonstrated the fidelity of this system with peptides of known KIR specificity bound to HLA-C*05:01. We then screened a peptide library to identify novel strong KIR2DS4 binding peptides presented by HLA-C*04:01. Peptide-exchanged HLA-C was functionally competent, promoting activation of KIR2DS4+ NK cells and inhibiting activation of KIR2DL1+ NK cells. Together, we show that peptide-exchangeable HLA-I molecules are ligands for KIR, presenting a flexible, efficient system for examining the peptide-sequence dependent recognition of HLA-I by KIR.

## Introduction

Natural killer (NK) cells are innate lymphoid cells that play a central role in host defence by eliminating virus-infected and malignant cells [1, 2]. NK cell responses are governed by integrating signals from activating and inhibitory receptors with a prominent role for class I human leukocyte antigen (HLA-I) binding receptors [3]. The killer cell immunoglobulin-like receptors (KIR) are a multigene family of activating and inhibitory receptors expressed on NK cells and T cells [4]. Many KIR have HLA-I ligands, and are associated with numerous human diseases including infection, cancer, autoimmunity and disorders of pregnancy [4, 5]. HLA-C is a ligand for multiple KIRs including KIR2DL1, KIR2DL2/3, KIR2DS1, KIR2DS2, KIR2DS4 and KIR2DS5 [6–12]. Where it has been examined, all HLA-I binding KIR exhibit peptide-dependent recognition of HLA-I with peptide positions 7 (p7) and p8 (of 9mers) proving most critical for receptor engagement [6, 13, 14]. Some KIR display limited peptide selectivity such as inhibitory KIR2DL1, while others display high levels of peptide specificity, such as activating KIR2DS4 [6, 13]. Recently, we proposed the ‘peptide-selectivity model’ where detection of HLA-I bound peptides by KIR plays a central role in understanding disease associations with specific combinations of KIR and HLA-I [15].

Despite the centrality of peptide specificity to certain KIR-HLA-I interactions, comprehensive mapping is limited by the practical challenges of screening panels of peptide–HLA-I (pHLA-I) complexes. Conventional approaches typically rely on cell lines engineered to express single HLA-I alleles with deficiency in transporter associated with antigen presentation (TAP) [8, 10]. These cell lines are time consuming to generate and clone-variable, presenting challenges for comparing KIR binding to different HLA-I allotypes. Likewise, producing soluble pHLA-I complexes by *in vitro* refolding one peptide at a time hinders library-scale screening, as even a modest screen of 50 peptides for a single HLA-I allotype could take months to generate [16].

To overcome these limitations, we established and validated two complementary peptide-exchange strategies: engineered “open MHC-I” that stabilizes HLA-I in a peptide-receptive state [17], and a dipeptide-mediated exchange method with MHC-I binding peptides modified at two positions that matter for KIR binding [18]. We combined these strategies with SpyTag-SpyCatcher chemistry to generate HLA-I that can be directly conjugated to plates or cells (CombiCells) [19] allowing orthogonal readouts such as ELISA and cell-based functional assays. Using both methods with HLA-C*05:01, we established the fidelity of peptide-exchange technologies for examining KIR binding, successfully reproducing known KIR-pHLA-I interactions [6]. Next, we established a peptide-exchange protocol for HLA-C*04:01 and screened a small peptide library to identify novel, functional ligands for the activating receptor KIR2DS4. Collectively, we report a scalable, generalizable platform for dissecting peptide-specific KIR recognition across HLA-I allotypes, with clear utility for mechanistic studies and translational applications.

## Results

### Overview and rationale for two peptide-exchange strategies

Previously, we and others generated TAP-deficient cells expressing single HLA-I alleles for the purpose of screening peptides for KIR binding [6, 8, 10, 11, 20, 21]. For each HLA-I allele, an independent TAP-deficient line is required (Figure 1A). Typically, HLA-I-null target cells (such as 721.221) are transduced with HLA-I allele of interest, followed by TAP disruption. Each step often requires flow sorting, drug selection and often clone screening, resulting in weeks to months per allele of interest. However, direct comparisons between different HLA-I allotypes are challenging as HLA-I expression levels cannot be precisely standardized across independently engineered lines. Alternatively, KIR binding can be measured to soluble peptide–HLA-I complexes generated by *in vitro* refolding (Fig 1B) [6, 9, 22]. For each HLA-I allotype, recombinant heavy chain (HC) and β2m are refolded in the presence of peptide and purified [16, 23]. This must be repeated for each peptide, thus even a modest screen of ~50 peptides for a single HLA-I allotype requires ~50 independent refolds and purifications, extending the timeline to months.

**Figure 1:**
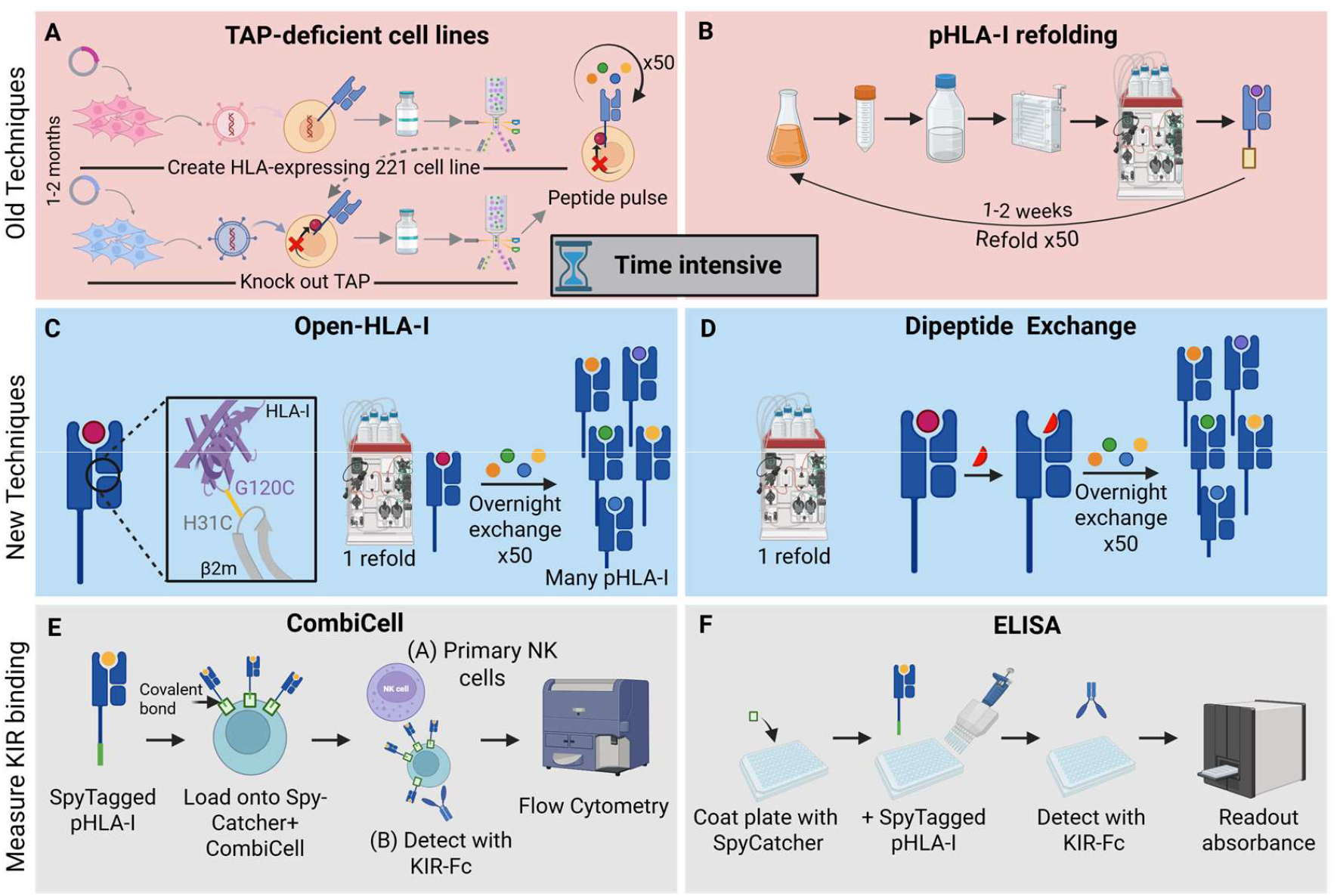
Strategies for interrogating KIR binding to pHLA-I. **(A)** Standard workflow to generate TAP-deficient HLA-I^+^ cell lines. HLA negative cells (721.221; 221) are transduced with lentivirus encoding HLA-I allele of interest. After selection and/or sorting for high expression, TAP is knocked out (via CRPISR/Cas9) or inhibited via delivery of TAP inhibitors such as ICP47. **(B)** Conventional soluble pHLA-I production by *in vitro* refolding. For each HLA-I allele, heavy chain (HC) is produced as bacterial inclusion bodies (1-2 weeks). Each peptide must be refolded individually (1-2 weeks). **(C)** “Open HLA-I” method utilises an inter-HC and β2m disulphide (G120C – H31C), which enables rapid placeholder-to-target peptide exchange from a single refold. **(D)** Dipeptide-mediated double exchange. HLA-I refolded with placeholder peptide is first exchanged with dipeptide, followed by exchange to target peptide. **(E)** Cell-based Combi readout: CHO cells expressing surface SpyCatcher capture SpyTag–pHLA-I for primary NK-cell functional assays or KIR-Fc binding. **(F)** Cell-free ELISA readout: SpyCatcher-coated plates covalently capture SpyTag–pHLA-I for KIR-Fc binding assays. TAP = Transporter associated with antigen presentation. (Created in http://Biorender.com)

A potential solution is to use ‘peptide-exchange’ technologies using refolded HLA-I molecules (Fig. 1B), which allow rapid replacement of a ‘placeholder’ peptide with the peptide of interest (Fig. 1C-F). Here, we tested two different methodologies of peptide exchange, peptide-receptive MHC-I “open MHC-I” and dipeptide-mediated exchange for peptides with amino acid substitutions at two positions [17, 18, 24]. Open MHC-I utilizes an engineered interchain disulfide across the HC/β2m interface that stabilizes an “open,” peptide-receptive MHC-I conformation with increased thermostability [17]. Cysteine substitutions at G120 (HC) and H31 (β2m), allow HC-β2m dimers to form that can be refolded with a placeholder peptide and efficiently exchanged with peptides of interest [17]. For dipeptide-mediated exchange, a two-step strategy is employed [18]. Unmodified HLA-I is refolded with a placeholder peptide, then challenged with a dipeptide to displace the placeholder peptide, and finally loaded with the peptide of interest [18, 24]. Dipeptides are typically common C-terminal residues for a specific HLA-I allotype bonded with an N terminal Gly such as GL, GM, and GV [18, 24]. For both methods, a single HLA-I refold is required, with peptide exchange performed via one or two overnight steps.

We combined these peptide-exchange technologies with SpyTag-SpyCatcher chemistry [25] to allow flexible KIR binding measurements across multiple formats. SpyTag and SpyCatcher act as ‘molecular glue’ such that when the 13 amino acid SpyTag interacts with SpyCatcher (138 residues, 15KDa), it forms a spontaneous iso-peptide bond in approximately 15 mins [25]. All peptide-HLA-I (pHLA-I) ligands were produced with a C-terminal SpyTag enabling covalent coupling to SpyCatcher absorbed directly on plates or expressed on the cell surface of CHO-K1 ‘CombiCells’ [19]. These CombiCells are engineered to lack ICAM-1 and display SpyCatcher using a short GPI-anchored hinge from human CD52 (Fig. 1E). Purified SpyCatcher was coated onto MaxiSorp plates to capture SpyTagged pHLA-I, followed by KIR–Fc conjugated to protein-A-HRP (Fig. 1F). Purified SpyCatcher was coated onto MaxiSorp plates to capture SpyTagged pHLA-I, followed by KIR–Fc conjugated to protein-A-HRP (Fig. 1F).

### Peptide exchange strategies for HLA-C*05:01

We evaluated KIR-Fc binding with peptide-exchanged open-HLA-C*05:01 (Fig. 2A&B) and after dipeptide-mediated exchange (Fig. 2C) [18]. For open-HLA-C*05:01, HLA-C*05:01(G120C) was refolded with β_2_m(H31C) and a “placeholder” peptide P2-EE (IIDKSG**<u>EE</u>**V) that does not support KIR binding, and is derived from ‘self’ peptide P2 (IIDKSGSTV) [6–8]. The placeholder was then exchanged with a panel of five peptide P2 (IIDKSG**<u>xx</u>**V) variants that differ only at p7 and p8, which are the principal KIR binding determining residues [6, 10, 13]. The panel included p7p8 combinations within peptide P2 of known specificities for KIR2DL1 (ST, QH, MW, IA, AV), KIR2DS1 (IA, AV) and KIR2DS4 (MW) [6].

**Figure 2:**
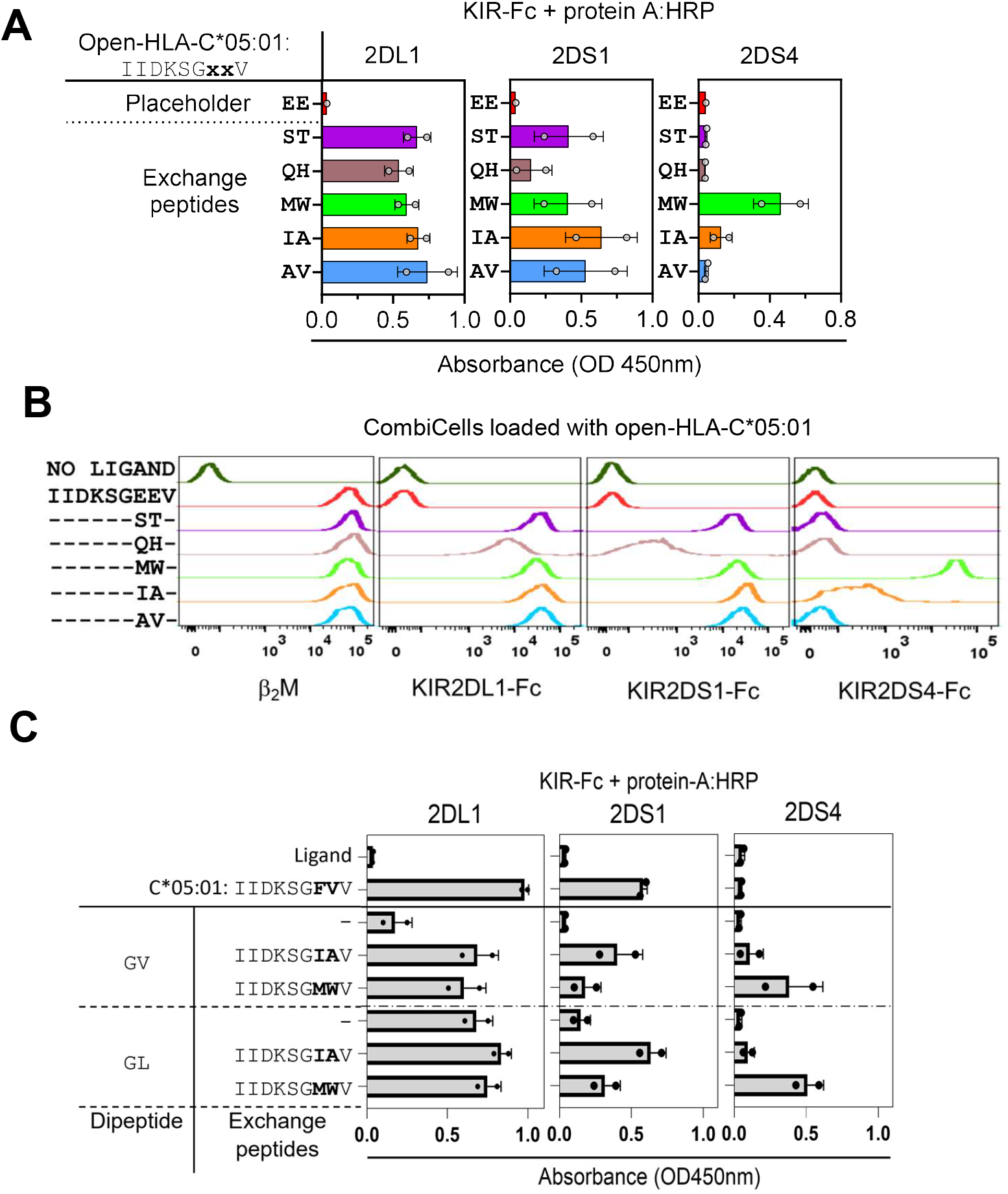
KIR binding to peptide-exchanged HLA-C*05:01. **(A**,**B)** KIR-Fc binding (KIR2DL1, KIR2DS1, KIR2DS4) binding to open-HLA-C*05:01 with placeholder peptide P2-EE or exchanged to peptides with indicated residues at p7p8. Open-HLA-C*05:01 was conjugated to plates in (A) and CombiCells in (B) via SpyCatcher. KIR-Fc binding was measured by ELISA in (A) and flow cytometry in (B), with level of conjugation displayed by anti-β2m mAb. **(C)** KIR-Fc binding to HLA-C*05:01(P2-FV) after dipeptide exchange (GV or GL), followed by exchange with peptides P2-MW or P2-IA. KIR-Fc binding was measured by ELISA.

As detected by ELISA, KIR2DL1-Fc recognized all five open-HLA-C*05:01 complexes after peptide-exchange, while un-exchanged open-HLA-C*05:01 containing P2-EE displayed no KIR2DL1-Fc binding, as expected (Fig. 2A). Consistent with previous studies, KIR2DS1-Fc exhibited a more selective profile, with the strongest binding to P2-IA and weaker binding to P2-QH [6]. KIR2DS4-Fc binding was highly restricted, showing robust recognition of P2-MW, a weak signal for P2-IA and no binding to ST, QH or AV (Fig. 2A). We next tested KIR-Fc binding to the same peptide-exchanged open-HLA-C*05:01 molecules displayed on CombiCells, measuring binding by flow cytometry. Unexchanged and peptide-exchanged open-HLA-C*05:01 showed robust staining with a β2m mAb, while KIR-Fc staining reflected known peptide sequence dependent binding patterns, mirroring the ELISA data and previous studies (Fig. 2B) [6]. In particular, KIR2DS4-Fc displayed robust staining to known KIR2DS4 binder P2-MW, weak binding to P2-IA and no binding to any other peptides.

We next tested the dipeptide-exchange method, where wild-type HLA-C*05:01 was refolded with the KIR2DL1-binding peptide P2-FV (IIDKSG**<u>FV</u>**V) and subjected to a two-step exchange: an initial incubation with GV or GL dipeptides, followed by exchange with full-length P2-IA or P2-MW. Exchange with dipeptide GV, lead to significant loss of KIR2DL1-Fc binding when compared to unexchanged HLA-C*05:01-P2-FV, consistent with P2-FV displacement. KIR2DS1-Fc, similar to KIR2DL1, bound the initial P2-FV complex, lost binding after dipeptide exchange, and regained binding upon reconstitution with P2-IA. Further, KIR2DS4-Fc showed negligible binding to the starting P2-FV complex but acquired robust binding after exchange to P2-MW. HLA-C*05:01-P2-FV also readily exchanged with dipeptide GL, with loss of KIR2DS1-Fc binding with GL alone, and gain of KIR2DS1 and KIR2DS4 binding after exchange to P2-IA and P2-MW, respectively. Intriguingly, despite efficient GL mediated peptide-exchange, KIR2DL1-Fc binding did not reduce in the presence of GL, compared to GV (Fig. 2C). It is unclear whether this is due to incomplete peptide exchange or if KIR2DL1 is able to bind HLA-C*05:01 loaded only with GL. Collectively, these results demonstrate that both the open-MHC-I and dipeptide-exchange protocols enable controlled, position-specific manipulation of the bound HLA-C*05:01 peptide and reproduced KIR-specific binding hierarchies. Also, we show that combining peptide exchange technologies with SpyTag-SpyCatcher chemistry allows for easy detection of KIR binding via ELISA or flow cytometry.

### Peptide-exchanged HLA-C*05:01 retains KIR2DL1-mediated inhibition comparable to wild-type

We next evaluated whether peptide-exchanged HLA-C*05:01 supports functional recognition by primary KIR2DL1+ NK cells comparable to wild-type HLA-C*05:01. To do so, we first established methods to activate NK cells using CombiCells as target cells. We generated recombinant ligands ULBP1 and CD155, which engage activating receptors NKG2D and DNAM-1, respectively [3]. Each ligand carried a C-terminal SpyTag allowing rapid conjugation to SpyCatcher expressed on CombiCells [19]. Ligand display was verified by antibody and receptor-Fc binding to Spytagged ligands loaded on CombiCells (Fig. 3A). We generated three formats of SpyTagged HLA-C*05:01: wild-type, open-MHC-I and dipeptide-exchanged HLA-C*05:01, each presenting strong KIR2DL1 ligand P2 (IIDKSGSTV) [8]. When mixed with primary NK cells, CombiCells displaying activating ligands CD155 and ULBP1 alone induced degranulation of both KIR2DL1+ and KIR2DL1-NK cells, measured by CD107a detection (Fig. 3B&C). CombiCells displaying CD155, ULBP1 and HLA-C*05:01 of any format lead to robust inhibition of KIR2DL1^+^, but not KIR2DL1^-^ NK cells (Fig. 3B&C). Thus, across both exchange strategies, peptide-exchanged HLA-C*05:01 complexes were indistinguishable from wild-type HLA-C*05:01 in their capacity to engage KIR2DL1 and completely inhibit NKG2D/DNAM-1–driven NK-cell activation.

**Figure 3:**
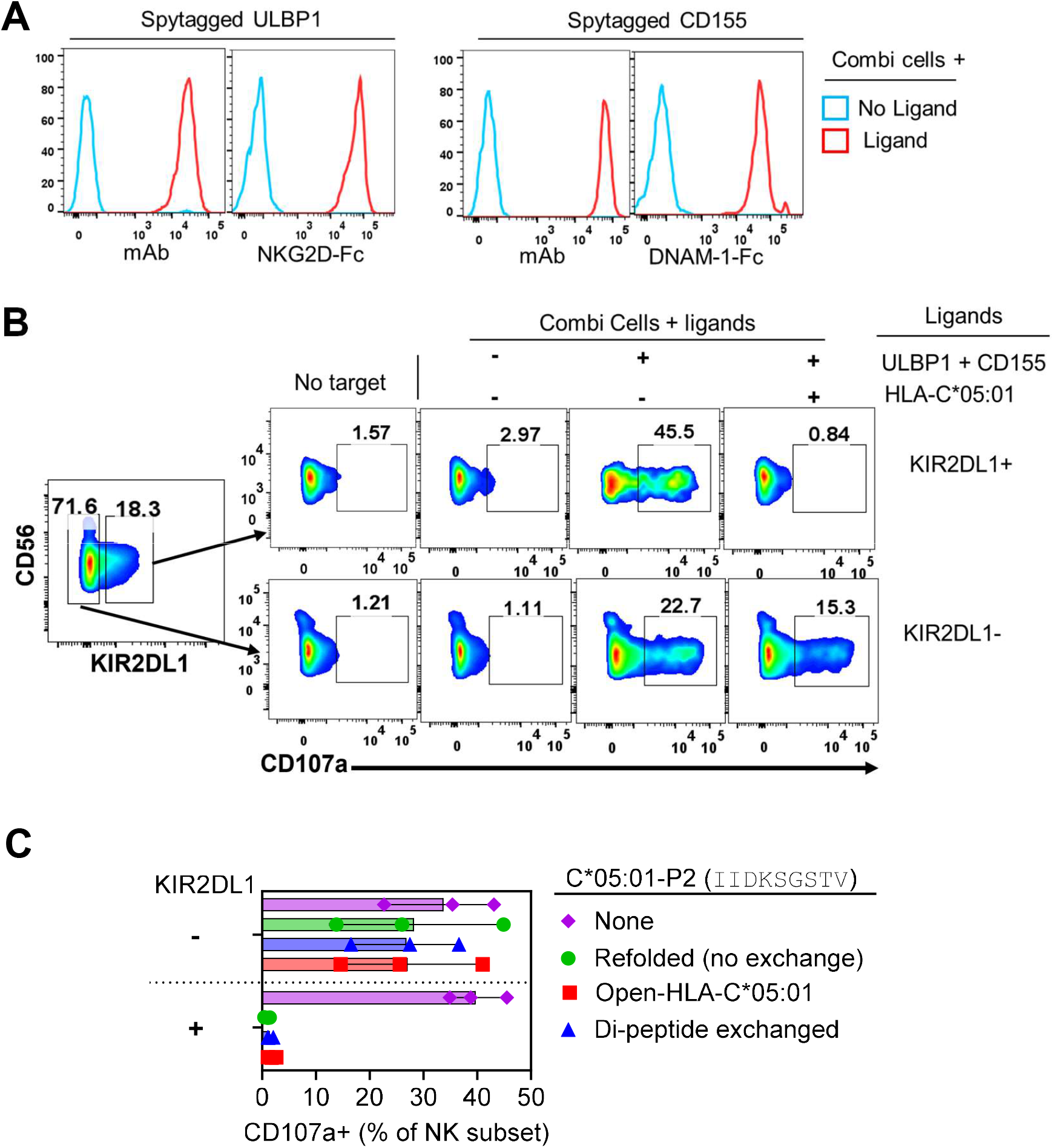
Recognition of peptide exchanged HLA-C*05:01 by primary KIR2DL1+ NK cells. **(A)** Display of ULBP1 and CD155 on CombiCells detected by mAbs and NKG2D-Fc, and DNAM-1-Fc, respectively. **(B&C)** KIR2DL1+ and KIR2DL1-NK-cell degranulation (CD107a upregulation) in response to ULBP1, CD155, and HLA-C*05:01 displayed on CombiCells. **(C)** Three types of HLA-C*05:01 were tested, all containing P2 (IIDKSGSTV); wild-type, open and dipeptide exchanged. Data from three independent experiments with NK cells from separate donors are shown.

### Developing a peptide exchange strategy for HLA-C*04:01

We previously identified multiple strong activating KIR binding peptides presented by HLA-C*05:01 [6, 7], but we lack similar ligands for other HLA-C allotypes. To establish these peptide-exchange platforms for screening peptide-libraries on a different HLA-C allotype, we focused on HLA-C*04:01. Previous studies identified the peptide QYDDAVYKL (#1) as a strong and weak ligand for KIR2DL1 and KIR2DS1, respectively [9, 10, 26], while the p8 Glu variant (#1-K8E; QYDDAVY**<u>E</u>**L) had no KIR binding. To create an exchangeable scaffold, we generated the open-MHC-I format of HLA-C*04:01 (open-HLA-C*04:01) [17]. We refolded HLA-C*04:01(G120C) heavy chains with β_2_m(H31C) (Fig. 4A) with either peptide #1 or #1-K8E. Both pHLA-I complexes eluted at ~56 mL by size-exclusion chromatography (Fig. 4A) and migrated as expected by SDS–PAGE generating a single band under non-reducing conditions, and two bands under reducing conditions (Fig. 4A). We then performed reciprocal peptide exchange between #1 and #1-K8E on open-HLA-C*04:01 and assessed peptide-exchange efficiency by measuring KIR2DL1-Fc binding by flow cytometry. As expected, open-HLA-C*04:01 refolded with #1 displayed strong KIR2DL1-Fc binding, while complexes refolded with #1-K8E displayed no binding (Fig. 4B). However, we observed no peptide exchange for both peptides, as KIR2DL1-Fc binding was unchanged post peptide-exchange (Fig. 4B).

**Figure 4:**
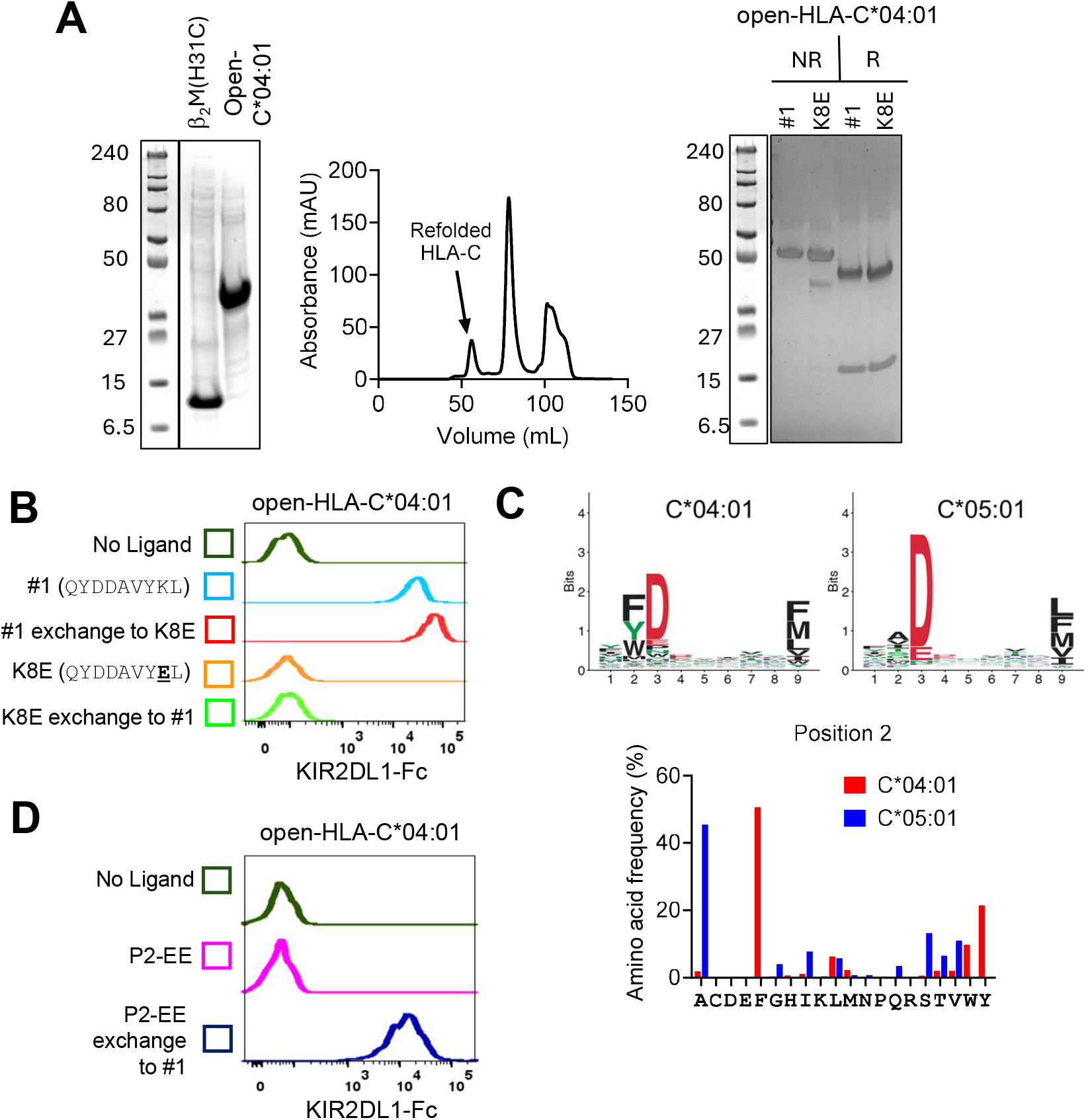
Development and validation of a peptide-exchange strategy for HLA-C*04:01. **(A)** SDS–PAGE of purified inclusion bodies (IBs) from open-C*04:01 heavy chain (G120C) and β2M (H31C (*left*). Size-exclusion chromatography (SEC) trace from purification of refolded open-HLA-C*04:01 complex (*middle*). SDS-PAGE of refolded open-HLA-C*04:01 complexes loaded with peptide #1 or K8E, run under non-reducing and reducing conditions (*right*). (**B)** KIR2DL1-Fc binding to open-HLA-C*04:01 displayed on CombiCells following indicated peptide exchange. **(C)** Peptide-binding motifs of HLA-C*04:01 and -C*05:01 (*top*). Amino acid frequency at position 2 for HLA-C*04:01 and HLA-C*05:01 (*bottom*). **(D)** KIR2DL1-Fc binding to open-HLA-C*04:01 displayed on CombiCells loaded with placeholder P2-EE or after exchange to #1. #1=QYDDAVYKL, K8E=QYDDAVY**E**L, P2-EE=IIDKSG**EE**V.

We reasoned that peptide-exchange failed as the placeholder peptide bound HLA-C*04:01 too strongly. HLA-C*05:01 and HLA-C*04:01 share similar peptide preferences at p3 for Asp and conserved hydrophobic residues at the C-terminus, but differ significantly at p2 with HLA-C*04:01 preferring large aromatic residues (Phe, Tyr and Trp) and HLA-C*05:01 preferring smaller residues such as Ala and Ser (Fig. 4C) [27, 28]. Therefore, we refolded open-HLA-C*04:01 with the HLA-C*05:01 binding peptide P2-EE (IIDKSGEEV) that lacks the aromatic p2 anchor commonly seen in HLA-C*04:01 binding peptides. This complex showed no KIR2DL1-Fc binding, but after exchange to #1 acquired robust KIR2DL1-Fc recognition (Fig. 4D), demonstrating successful peptide-exchange. Thus, peptide exchange with open-HLA-C*04:01 is feasible by using a suboptimal binding placeholder peptide.

### Identification of functional KIR2DS4 binding peptides via screening with open-HLA-C*04:01

In previous work, the first KIR2DS4 binding peptide we identified presented by HLA-C*05:01 carried Tyr at p8, P2-AY [7]. Through targeted substitutions and screening peptide-libraries, we identified multiple additional KIR2DS4 binding peptides, such as P2-MW [6, 7]. After the discovery of P2-AY, in unpublished work (shown now here), we also tested four p8 Tyr containing peptides from the HLA-C*04:01 immunopeptidome [29] for KIR2DS4 binding using a previously developed HLA-C*04:01+ TAP deficient cell line [30]. We identified ‘self’ peptides VYDVRQAYV (named VV9 here), and AYDDKIYYF (AF9) as a strong and intermediate KIR2DS4 binding peptides, respectively (Fig. 5A). VV9 is derived from residues 191-199 of URB1, nucleolar pre-ribosomal-associated protein 1 (Uniprot ID: O60287). Both peptides also stabilised HLA-C*04:01 and bound KIR2DL1 (Fig. 5A).

**Figure 5.**
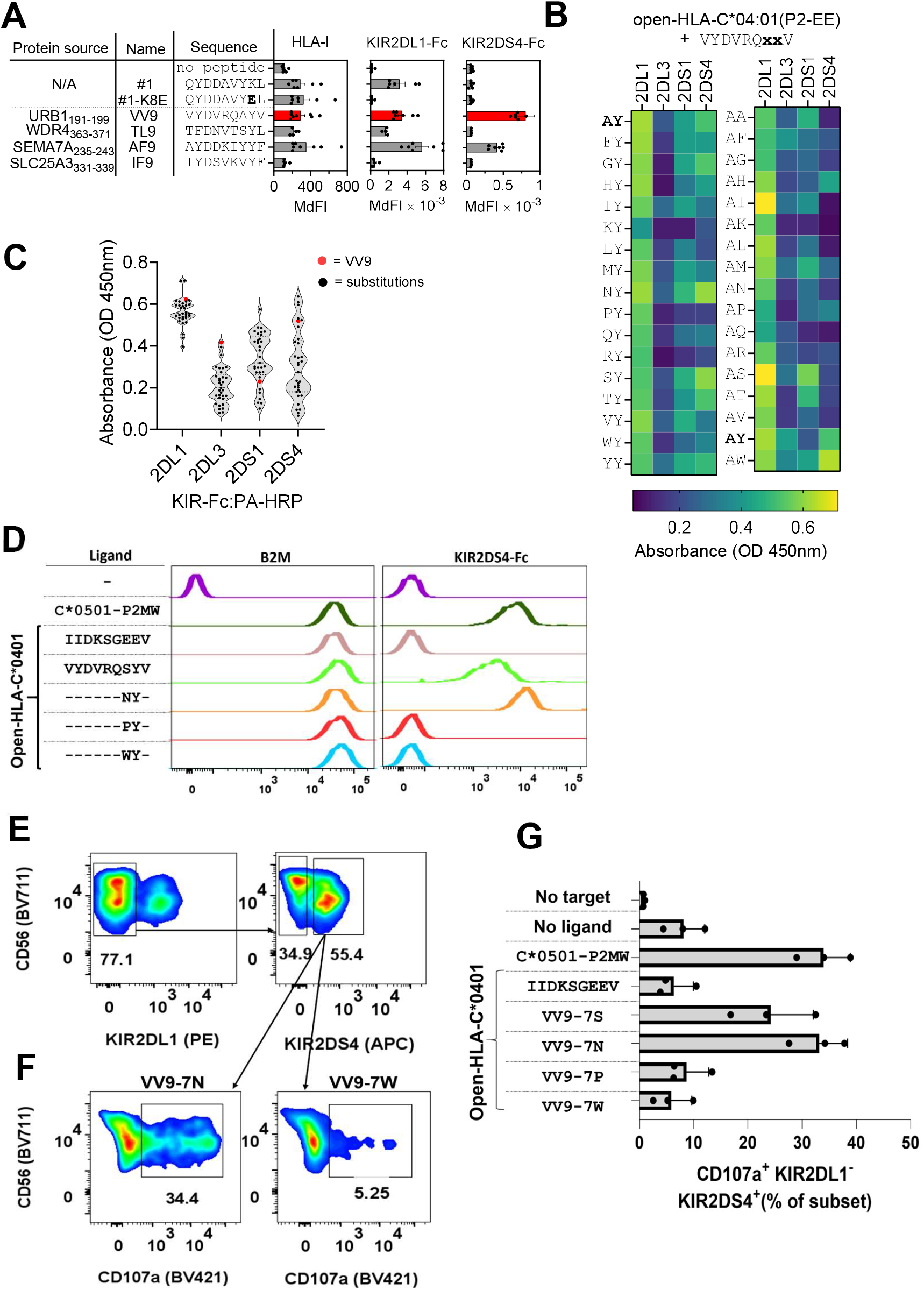
Screening and functional validation of KIR2DS4-binding peptides with open-HLA-C*04:01. **(A)** KIR2DL1-Fc, KIR2DS4-Fc and HLA-I mAb binding to 221-C*04:01-ICP47 cells preloaded with peptides #1, K8E, and four ‘self’ peptides containing p8 Tyr. Source proteins for the ‘self’ peptides are shown. **(B)** KIR-Fc (KIR2DL1, KIR2DL3, KIR2DS1, KIR2DS4) binding to open-HLA-C*04:01 exchanged with VV9 p7 (VYDVRQ**x**YV) and p8 (VYDVRQA**x**V) library peptides. Binding was measured by ELISA. **(C)** Data from (B) shown as violin plots with VV9 (red) and VV9-substituted ligands (black). **(D)** KIR2DS4-Fc and β2M antibody binding to open-HLA-C*04:01 displayed on CombiCells and exchanged with selected VV9 peptides. **(E&F)** Flow-cytometric gating strategy used to identify the KIR2DS4^+^ KIR2DL1-NK cell population and response to CombiCells displaying open-HLA-C*04:01 loaded with VV9-7S or VV9-7W (F). **(G)** Degranulation of KIR2DS4^+^ KIR2DL1^-^ cells as in F, following exposure to CombiCells displaying HLA-C*05:01-P2-MW or open-HLA-C*04:01 loaded with indicated peptides. Data from three independent experiments using NK cells from three separate donors are shown.

To explore the potential of peptide-exchange systems for library screening, we used open-HLA-C*04:01 refolded with P2-EE (Fig. 4) and tested a peptide library based on VV9. Two complementary libraries were synthesized with either fixed positions p7 (A) or p8 (Y) and every other amino acid (excluding Cys, Asp and Glu) were tested at p8 or p7, respectively. There were 33 peptides in total with the format VYDVRQ**x**YV (p7 library) or VYDVRQA**x**V (p8 library). Each peptide was exchanged onto Spytagged open-HLA-C*04:01, then captured on SpyCatcher-coated plates before detection of KIR-Fc binding by ELISA. We tested KIR2DL1-Fc, KIR2DL3-Fc, KIR2DS1-Fc, and KIR2DS4-Fc binding across the peptide panel (Fig. 5B&C). KIR2DL1-Fc bound broadly to most variants with strongest binding to VV9-8S (VYDVRQA**<u>S</u>**V) [8, 10, 13]. We previously showed that cross-reactive binding of KIR2DL3 to C2-HLA-C allotypes like HLA-C*04:01 is dependent on peptide sequence [6, 8]. However, none of the VV9 derived peptides conferred strong binding of KIR2DL3-Fc to HLA-C*04:01, with the strongest binding conferred by VV9 itself (Fig. 5B&C). For KIR2DS1 most substitutions improved binding, with the strongest binding to VV9-8S, the same as KIR2DL1 (Fig. 5B&C). Several substitutions particularly those with positive charges at p7 and p8 (AK, AR, KY and RY) diminished KIR2DS1 but not KIR2DL1 binding, consistent with KIR2DS1 being a more peptide-selective receptor than KIR2DL1 [6]. For KIR2DS4, three substitutions increased binding compared to VV9 and they were; VV9-8W (VYDVRQA**<u>W</u>**V), VV9-7N (VYDVRQ**<u>N</u>**YV) and VV9-7S (VYDVRQ**<u>S</u>**YV). Three additional substitutions; VV9-7T (VYDVRQ**<u>T</u>**YV), VV9-7G (VYDVRQ**<u>G</u>**YV) and VV9-7Y (VYDVRQ**<u>N</u>**YV) had minimal impact while the remaining substitutions decreased KIR2DS4 binding to a greater or lesser extent. This is consistent with the high peptide specificity of KIR2DS4 as the interaction is highly intolerant to amino acid substitutions, in sharp contrast to KIR2DL1 (Fig. 5B&C).

We next tested whether these newly identified VV9 variants can activate primary KIR2DS4^+^ NK cells. Using open-HLA-C*04:01 loaded on CombiCells VV9-7N displayed stronger KIR2DS4 binding than VV9-7S, with VV9-7N displaying similar binding strength to HLA-C*05:01-P2-MW (Fig. 5D). In contrast VV9-7P and VV9-7W displayed no KIR2DS4-Fc binding (Fig. 5D). We next tested whether these complexes could induce functional activation of KIR2DS4^+^ NK cells. Primary NK cells from KIR2DS4^+^ donors were mixed with CombiCells loaded with HLA-C*05:01-P2-MW or open-HLA-C*04:01 exchanged with VV9 variants. We gated on KIR2DS4^+^ KIR2DL1^-^ NK cells to exclude confounding inhibition via their shared ligand specificity (Fig. 5E&F). Consistent with KIR2DS4 binding data, both VV9-7N and VV9-7S elicited clear activation of KIR2DS4^+^ NK cells, while the non-binder peptides (VV9-7P and VV9-7W) failed to activate KIR2DS4^+^ NK cells (Fig. 5G). We observed similar levels of KIR2DS4^+^ NK cell activation with open-HLA-C*04:01-VV9-7N and HLA-C*05:01-P2-MW, suggesting VV9-7N is a similarly strong KIR2DS4 ligand [6]. Thus, screening peptide libraries with open-HLA-C*04:01 is a feasible approach to identify novel, strong, functional peptide binders for KIR2DS4. Together, our data show that peptide-exchange technologies are likely to be useful tools for identifying strong KIR binding peptides.

## Discussion

KIR binding to HLA-I is peptide sequence dependent, and for several KIR this dependency rises to striking peptide specificity [13]. Yet, practical dissection of this specificity is limited by the available tools to examine KIR binding in the presence of defined peptide sequences. Deriving TAP-deficient cells expressing single HLA-I alleles is labour-intensive, and these cell lines display clone-to-clone heterogeneity with variable capacities for peptide presentation. Indeed, HLA-I alleles display variable resistance to TAP knockout or inhibition [31–33], complicating the comparison of KIR binding to different alleles. In contrast, recombinant pHLA-I production offers tight control of peptide sequence via *de novo* refolding but lacks scalability for screening peptide libraries. These constraints urged us to explore alternative approaches that can increase the throughput and consistency of assays designed to interrogate KIR peptide-specificity.

Here, we combined peptide-exchange strategies with SpyCatcher–SpyTag chemistry [25] to develop a streamlined, generalisable workflow to assemble defined pHLA-I complexes, map KIR specificity, and functionally validate KIR-pHLA-I interactions. We used open-HLA-C*05:01 refolded with a non-KIR-binding placeholder peptide (P2-EE) such that successful exchange was detected by gain of KIR binding. This approach faithfully reproduced known KIR binding specificities and hierarchies for five P2 p7p8 variants establishing a robust proof of concept for the methodology. For the dipeptide method, we tested two dipeptides, GV and GL based on F-pocket preferences of HLA-C*05:01 [8, 27] and adopted a sequential exchange strategy such that dipeptide displacement was detected by loss of KIR binding, followed by gain of binding upon loading of a new full-length peptide. For both dipeptides, subsequent exchange with full-length peptides restored the expected receptor specificities, confirming that the dipeptide intermediate was replaced. However, the expected loss of KIR2DL1-Fc binding upon dipeptide exchange was only observed with GV and not GL. This does not appear to be due to lack of GL displacement of the placeholder peptide P2-FV, as KIR2DS1-Fc binding dramatically reduced with GL alone, and exchange with P2-MW conferred binding to KIR2DS4-Fc. A potential explanation is that GL bound to HLA-C*05:01 is sufficient for KIR2DL1 binding. This is consistent with the idea that KIR2DL1 has minimal peptide preferences when binding HLA-C*05:01, including P2-GG that contains no sides chains at p7p8 [6, 8]. For applications where strict native conformation is essential, this wild-type dipeptide sequential approach is preferable to engineered conditional systems such as open-HLA-I.

While we have begun to identify KIR binding peptides across some HLA-C allotypes [6], many allotypes have not been studied. It is also unclear whether the same sequence ‘rules’ that confer KIR binding on one HLA-I allotype will apply to all. We therefore turned to HLA-C*04:01, a common HLA-C allotype across many populations [34]. We first established that peptide-exchange could occur with open-HLA-C*04:01. To our surprise, open-HLA-C*04:01 refolded with #1 or #1-K8E failed to display any measurable peptide-exchange. However, when refolded with P2-EE, we observed peptide-exchange with #1 and gain of KIR2DL1-Fc binding. P2-EE is a strong HLA-C*05:01 binding peptide with a suboptimal p2 anchor for HLA-C*04:01 that was sufficient for refolding and peptide-exchange, but yielded less material than HLA-C*04:01 refolds with high-affinity peptides. Notably, open-HLA-C*05:01 tolerated exchange with a high-affinity placeholder peptide, whereas HLA-C*04:01 did not, suggesting allotype-specific differences. Practically, optimizing exchange will require empirical selection of placeholder peptides spanning a range of affinities to balance refolding yield with exchange efficiency.

After establishing this peptide-exchange method for HLA-C*04:01, we determined KIR binding to a library of 33 peptide variants of ‘self’ peptide VV9 with p7 or p8 substitutions. Consistent with previous work, KIR2DL1 binding to HLA-C*04:01 was largely resistant to amino acid substitutions at p7 and p8 [6, 8]. This is in sharp contrast to the activating KIR that display greater peptide-specificity [6] and were therefore more susceptible to changes in p7p8 sequence. A few substitutions increased KIR2DS4 binding to VV9, including p8 Trp found in many KIR2DS4 binding peptides presented by HLA-C*05:01, HLA-C*08:02 and HLA-C*16:02 [6, 7]. However, presence of p8 Trp or Tyr is not sufficient to confer KIR2DS4 binding as most p7 substitutions reduced KIR2DS4 binding, consistent with previous data [6, 7]. Much like P2-AW and HLA-C*05:01 [6], p7 sequence modulates KIR2DS4 binding in the presence of a p8 Tyr/Trp. We validated these novel KIR2DS4 binders with primary KIR2DS4+ NK cells, demonstrating the full utility of our system to identify novel functional ligands.

The advantages of our system are speed, flexibility, scalability and cost. Refolding HLA-I molecules is quick (days) and uses established methods refined over the past 30 years [16,23]. While each allotype may require some optimisation as we found for HLA-C*04:01, we quickly arrived at a rational solution following a biochemical understanding of HLA-I peptide binding (using a suboptimal binding placeholder peptide). Further, the flexibility of the CombiCell system [19] allows for rapid validation of KIR binding in functional experiments that can be modified be to study NK cell activation or inhibition. Currently, our lab is using the CombiCells to explore how NK cells integrate signals from multiple receptors concomitantly across a range of ligand densities. The method is highly scalable. It required only two 96 well plates to determine the binding of four KIRs to 33 peptides presented by HLA-C*04:01. A larger library or more receptors could be incorporated with little difficulty, perhaps via 384 well plates. However, the biggest barrier to larger screens remains the cost of synthetic peptide libraries, with the cheapest peptides costing approximately £15-25 each and thus screening hundreds or thousands of sequences would be prohibitively expensive.

Our approach should yield low false positive rates as KIR binding requires both efficient peptide exchange and KIR engagement. However, this method is susceptible to false negatives as they could arise from inefficient peptide-exchange, poor KIR binding or both. This is one advantage of TAP-deficient cell systems where peptide-stabilisation is relatively easy to measure. However, examining peptide binding affinities to HLA-C [35] alongside KIR binding assays could be a complementary strategy to rule out inefficient peptide-exchange as an explanation for negative results. While not tested here, our approach would likely be equally successful with UV labile peptides [36].

Together, we present an efficient, versatile platform for peptide exchange that supports precise interrogation of peptide-specific KIR binding to HLA-I, which can also be extended to other HLA-I binding receptors. These tools should accelerate discovery of novel KIR binding peptides providing insight into basic KIR function and their roles in human diseases [15].

## Materials and methods

### Recombinant proteins

Plasmids (pET-30a) encoding the genes for recombinant HLA-C*05:01 and HLA-C*04:01 heavy chains with a C-terminal SpyTagv3 (RGVPHIVMVDAYKRYK) and human β2-microglobulin (β2m) including the G120C (heavy chain) and H31C (β2m) variants were expressed in *E. coli* BL21(DE3) as inclusion bodies and purified as described [16]. For refolding, 15 mg heavy chain, 7 mg β2m, and 5 mg peptide (GenScript) were combined in 0.5 L refolding buffer (3 M urea (Merck, Cat. No: U1250-5KG), 0.4 M L-arginine-HCl (Thermo Scientific, Cat. No: J11500.A1), 0.1 M Tris-HCl pH 8.0 (Thermo Scientific, Cat No: J22638.AE), 2 mM Na-EDTA (Invitrogen, Cat. No: 15575020), 0.16% (w/v) reduced glutathione (Thermo Scientific, Cat No: J62166.22), 0.03% (w/v) oxidized glutathione (Thermo Scientific, Cat. No: 320225000) and stirred at 4 °C for 48 h. The mixture was dialyzed at 4 °C against 10 L of 10 mM Tris-HCl pH 8.0 with three buffer changes over 12 h, and concentrated using a Vivaflow device (10 kDa MWCO cut-off). One ml of concentrate was purified by size-exclusion chromatography on a HiLoad® 16/600 Superdex® 75 pg column. Purified proteins were further confirmed in reduced and non-reduced conditions using sodium dodecyl sulphate-polyacrylamide (SDS-PAGE) gel electrophoresis. Purified pHLA-I complexes were stored at −80°C until use.

Plasmids encoding the ectodomains of ULBP1 (pD649-HAsp-ULBP1-Fc(DAPA)-AviTag-6xHis) and CD155 (pMP71-hCD155) were obtained from Addgene. pMP71-hCD155 was a gift from Sébastien Walchli (Addgene plasmid # 118630; http://n2t.net/addgene:118630; RRID:Addgene_118630) [37]. pD649-HAsp-ULBP1-Fc(DAPA)-AviTag-6xHis was a gift from Chris Garcia (Addgene plasmid # 156597; http://n2t.net/addgene:156597; RRID:Addgene_156597) [38]. Ectodomains were cloned by PCR into pD649 each fused to a C-terminal SpyTagv3,was transfected into Expi293 cells using ExpiFectamine 293 (Gibco, Cat. No: A14524) and cultured for seven days at 37°C and 8% CO_2_. Culture supernatants were collected by centrifugation at 3,000 × g for 30 min at 4°C, and the proteins were purified by HisTrap affinity chromatography followed by size-exclusion chromatography (SEC). Purified proteins were stored in −80°C until further use.

### Binding of Fc fusion or β2M mAb on CombiCells

CombiCells (Chinese Hamster Ovarian (CHO-K1) cells with surface-displayed SpyCatcher and ICAM-1 knockout) were seeded at 5 × 10^4^ cells per well in flat-bottom 96-well plates and cultured overnight at 37 °C. SpyTag-tagged pHLA-I complexes or ligand (ULBP1 or CD155) were diluted to 0.5 μM in complete DMEM (10% FBS, 1% penicillin–streptomycin). The medium was aspirated, 50 μL of ligand was added per well, and cells were incubated for 40 min at 37 °C. Cells were washed twice with complete DMEM, detached by incubation in 10 mM Tris (pH 8.0) for 5 min at 37 °C, transferred to U-bottom plates, and washed with PBS (5 min, 300 × g). For binding assays, APC-conjugated Fc fusion protein (NKG2D-Fc (R&D Systems, Cat. No: 1299-NK) or DNAM-1-Fc (R&D Systems, Cat. No: # 666-DN) or KIR-Fc (R&D Systems, KIR2DL1-Fc (Cat. No: 1844-KR-050), KIR2DL3-Fc (Cat. No: 2014-KR-050), KIR2DS4-Fc (Cat. No: 1847-KR-050) (3.6 μg mL^-1^) was added at 25 μL per well and incubated for 30 min at 4 °C in the dark. For ligand detection, anti-β2M antibody (BioLegend, Cat. No:316306) or anti-CD155 (BioLegend, Cat. No: 337610) or anti-ULBP-1 (R&D Systems, Cat. No: FAB1380P) (1:75) was added at 25 μL per well. Cells were washed with PBS (5 min, 300 × g), resuspended in 150 μL PBS, and analysed by flow cytometry.

### Peptide exchange

Fifty μL of purified open-pHLA-I (G120C/H31C) complex [1μM] were incubated with peptide of interest at a 10-fold molar excess (pHLA-I:peptide = 1:10) overnight at room temperature (RT). For the dipeptide exchange, 50 μL of wild-type pHLA-I-P2FV [1μM] was first incubated with the dipeptide (glycyl-valine GV or glycyl-leucine GL) at a 20-fold molar excess (1:20) overnight at RT. The dipeptide-loaded complex was then exchanged by adding the full-length peptide at a 10-fold molar excess and incubating overnight at RT. Exchanged material was stored at −20 °C until use.

### ELISA

MaxiSorp Nunc-Immuno 96-well plates (Thermo Scientific, Cat. No:442404) were coated with 50 μL per well of SpyCatcher3 (BioRad, Cat. No: TZC025) (0.5 μM) and incubated overnight at 4 °C. The next day, wells were washed twice with PBS and blocked with 360 μL of 4% (w/v) skimmed milk in PBST (PBS + 0.05% Tween-20) for 1.5 h at room temperature (RT). After three PBST washes, 50 μL of SpyTag-tagged pHLA-I (0.5 μM in blocking buffer) was added to each well and incubated for 1 h at RT, followed by three additional PBST washes. KIR-Fc was diluted to 2 μg/mL in blocking buffer, 50 μL was added per well, and plates were incubated for 1 h at RT. Plates were washed three times with PBST, incubated with Protein A–HRP (Invitrogen, Cat. No: 101023) (1:10,000 in blocking buffer) for 1 h in the dark, washed again three times, and developed with 50 μL 1-Step Turbo TMB (Thermo Scientific, Cat. No: 34022) for 2.5 min. Reactions were stopped with 50 μL 1 M HCl (Merck, Cat. No:1090601000), and absorbance was measured at 450 nm on FLUOstar Omega micro plate reader (BMG LABTECH).

### Degranulation assay with primary NK cells

Peripheral blood mononuclear cells (PBMCs) were collected from leukocyte cones of healthy donors with informed consent and institutional approval (R83151/RE001, MS iDREC, University of Oxford) and isolated by Ficoll-Paque (Cytiva, Cat. No: GE17-1440-02) density-gradient centrifugation. NK cells were enriched using the StemCell Technologies NK Cell Isolation Kit (Cat. No: 17955) according to the manufacturer’s protocol and cultured in NK MACS medium (Miltyeni Biotec, Cat. No: 130-112-968) supplemented with IL-2 (500 IU/mL) and IL-15 (10ng/mL). For studying inhibition of KIR2DL1+ NK cells, CombiCells were incubated with 25 μL SpyTag-tagged pHLA-I (0.5 μM in complete DMEM) and 25 μL of each activating SpyTag-tagged ligand (ULBP1 and CD155; 0.1 μM each) for 40 min, then washed twice with complete DMEM. For activation of KIR2DS4+ NK cells, CombiCells were incubated with 50 μL SpyTag-tagged pHLA-I (0.5 μM in complete DMEM) for 40 min and washed twice. Primary NK cells were then added at an effector-to-target (E:T) ratio of 2:1 with anti-CD107a (BioLegend, Cat. No: 328626) at 1:100, plates were centrifuged at 300 × g for 1 min to promote contact with adherent targets, and incubated at 37°C for 3 h. NK cells were subsequently transferred to 96-well U-bottom plates, washed with PBS (5 min, 300 × g), and stained for 20 min at 4°C in the dark with mAbs to CD56 (BioLegend, Cat. No: 318336) (1:50), KIR2DL1 (R&D Systems, Cat. No: MAB1844) (1:50) and KIR2DS4 (Miltenyi Biotec, Cat. No: 130-092-0681) (1:50), followed by PBS washes. All flow cytometry data were acquired on BD LSRFortessa™ X-20 Cell Analyzer.

## Funding

This work was primarily supported by an MRC Career Development Award (MR/X020746/1) to MJWS. Additional support was provided by MSD Pump Priming award to MJWS, a Medical Sciences Graduate School Studentship award to BL, an NIH-OxCam Studentship awarded to EH, Wellcome Trust Grant 301534/Z/23/Z to OD.

## Author contributions

Conceptualisation: MJWS

Methodology: TMM, BL, OD, MJWS

Experiments: TMM, BL, EH, LC, MJWS

Funding acquisition: EL, OD, TE, MJWS

Supervision: MJWS, EL, TE

Writing – original draft: TMM, MJWS

Writing – review & editing: All

## Notes

### Competing Interest Statement

The authors have declared no competing interest.

